# An in vivo CRISPR screening platform for prioritizing therapeutic targets in AML

**DOI:** 10.1101/2020.12.28.424340

**Authors:** Shan Lin, Clément Larrue, Bo Kyung A. Seong, Neekesh V. Dharia, Miljan Kuljanin, Caroline Wechsler, Guillaume Kugener, Amanda Robichaud, Amy Conway, Thelma N. Mashaka, Sarah Mouche, Biniam Adane, Jeremy Ryan, Joseph D. Mancias, Scott T. Younger, Federica Piccioni, Lynn Lee, Mark Wunderlich, Anthony Letai, Jérôme Tamburini, Kimberly Stegmaier

## Abstract

CRISPR-Cas9-based genetic screens have successfully identified cell type-dependent liabilities in cancers, including acute myeloid leukemia (AML), a devastating hematologic malignancy with poor overall survival. Because most of these screens have been performed in vitro, evaluating the physiological relevance of these targets is critical. We have established a CRISPR screening approach using orthotopic xenograft models to prioritize AML-enriched dependencies in vivo, complemented by the validation in CRISPR-competent AML patient-derived xenograft (PDX) models tractable for genome editing. Our integrated pipeline has revealed several targets with translational value, including *SLC5A3* as a metabolic vulnerability for AML addicted to exogenous myo-inositol and *MARCH5* as a critical guardian to prevent apoptosis in AML. MARCH5 repression enhanced the efficacy of BCL2 inhibitors such as venetoclax, highlighting the clinical potential of targeting *MARCH5* in AML. Our study provides a valuable strategy for discovery and prioritization of new candidate AML therapeutic targets.

**Statement of significance:** There is an unmet need to improve the clinical outcome of AML. We developed an integrated in vivo screening approach to prioritize and validate AML dependencies with high translational potential. We identified *SLC5A3* as a metabolic vulnerability and *MARCH5* as a critical apoptosis regulator in AML, representing novel therapeutic opportunities.

## Introduction

AML is a heterogeneous hematologic malignancy characterized by the accumulation of abnormal myeloblasts. Despite the efficacy of chemotherapy and stem cell transplantation for some patients, cure rates for AML remain between 35-40% overall and less than 15% for older adults(1). Continued efforts are needed to identify new therapeutic approaches for these patients.

The successful adaptation of CRISPR-Cas9 approaches for genetic screens has become a powerful tool for the unbiased discovery of essential genes in mammalian cells(2,3). First-generation, large-scale functional genomic screens to identify the critical genes involved in cancer cell maintenance have been completed, such as the Broad Institute’s and Sanger Center’s Cancer Dependency Maps (Depmap, https://depmap.org/)(4,5). These efforts have revealed hundreds of potential genetic vulnerabilities in AML cells in vitro. However, to better leverage these data for guiding the development of new anticancer treatments, it is necessary to prioritize these gene targets to identify candidates with the best translational potential.

The niche may influence the physiological behavior of cancer cells, such as proliferation and response to therapy. Because large-scale genetic screens were primarily performed in vitro, an important next step in prioritizing new therapeutic targets is to validate their essentiality in a proper in vivo microenvironment. Human AML orthotopic disease modeling is highly physiologically relevant, as AML cells will engraft in the bone marrow microenvironment in the mouse. In vivo CRISPR screening has been performed to identify tumor suppressors, oncogenes and fitness genes in various cancer contexts(6,7), including several genetically engineered mouse models of hematologic malignancies (8–10). However, such an application in human AML orthotopic xenograft models has not been performed. Therefore, we optimized a protocol for CRISPR screening in orthotopic xenograft models to systematically evaluate the physiological relevance of top AML dependencies emerging from genome-scale CRISPR-Cas9 in vitro screens.

Although established AML cell lines are amenable for genetic screens, they cannot fully recapitulate all pathophysiological aspects of the disease. Target validation directly in primary patient samples is desirable, yet not readily accessible. Instead, the AML PDX has emerged as a valuable preclinical model that largely reflects the molecular and phenotypic characteristics of the primary disease(11,12). Thus, we developed PDX models amenable to genome-editing as a system for ensuring the translational relevance of the identified targets. By combining in vivo screening and CRISPR-competent PDX models, we devised an integrated pipeline to prioritize AML dependencies and investigated the top novel targets emerging from this approach.

## Results

### In vivo CRISPR-screens using xenograft models of human AML

To identify AML-enriched dependency genes, we explored the DepMap Avana CRISPR-Cas9 screen dataset and selected the gene targets that AML cells depend on for growth compared to the other cancer types included in the screen. This gene set was intersected with additional AML in vitro screen datasets including the combined Broad Institute and Novartis shRNA screens and a focused in vitro CRISPR screen in AML cell lines (13,14). These 200 top-ranked AML-enriched gene dependencies were enriched for various biological pathways, such as chromatin and transcription regulation, metabolic processes and mitochondria organization (**Figure S1A-B**). To distinguish the on-target from off-target anti-proliferative effects caused by CRISPR-mediated DNA cutting in regions of amplification (15,16), we designed three targeting single guide RNAs (sgRNAs) and three intronic control sgRNAs for each gene. With 120 additional negative control sgRNAs, a focused library with 1320 sgRNAs was constructed (**Figure 1A**).

**Figure 1.**
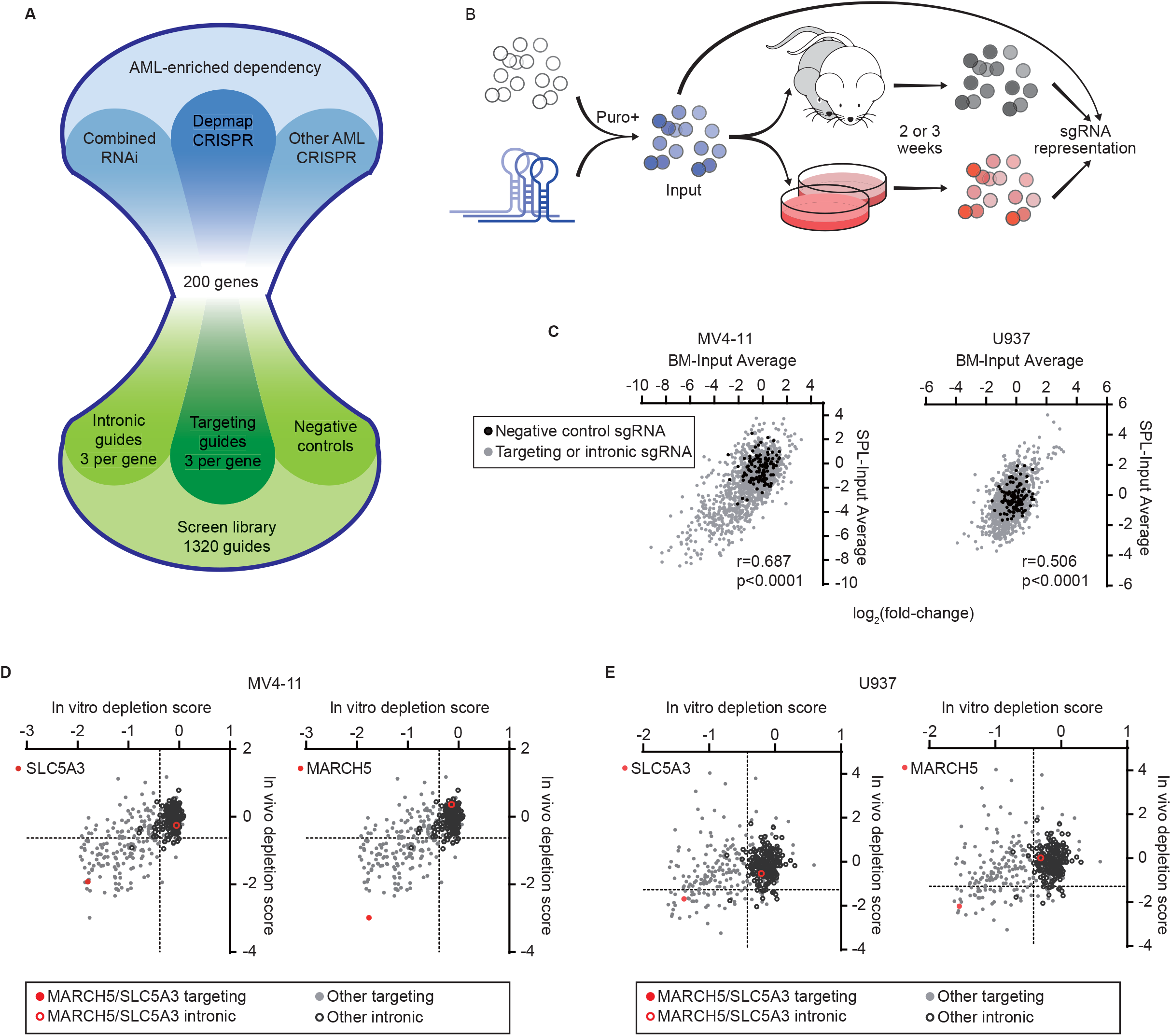
In vivo CRISPR screens prioritize genetic dependencies in human AML. **A.** Schematic of the library design. **B.** Schematic of in vivo and in vitro CRISPR screening approach. **C.** Scatterplot showing the correlation of relative abundance of sgRNAs in bone marrow (BM) versus spleen (SPL). Data points representing negative control sgRNAs (black solid circles) are indicated. **D, E.** Scatterplots showing the in vivo and in vitro depletion scores of *SLC5A3* and *MARCH5* at a gene level in MV4-11 (**D**) and U937 (**E**). Data points representing the median value of each intronic sgRNA set are indicated (black hollow circles). Scores of *SLC5A3* and *MARCH5* (red solid circle) and their intronic controls (red hollow circle) are highlighted.

**Figure 2.**
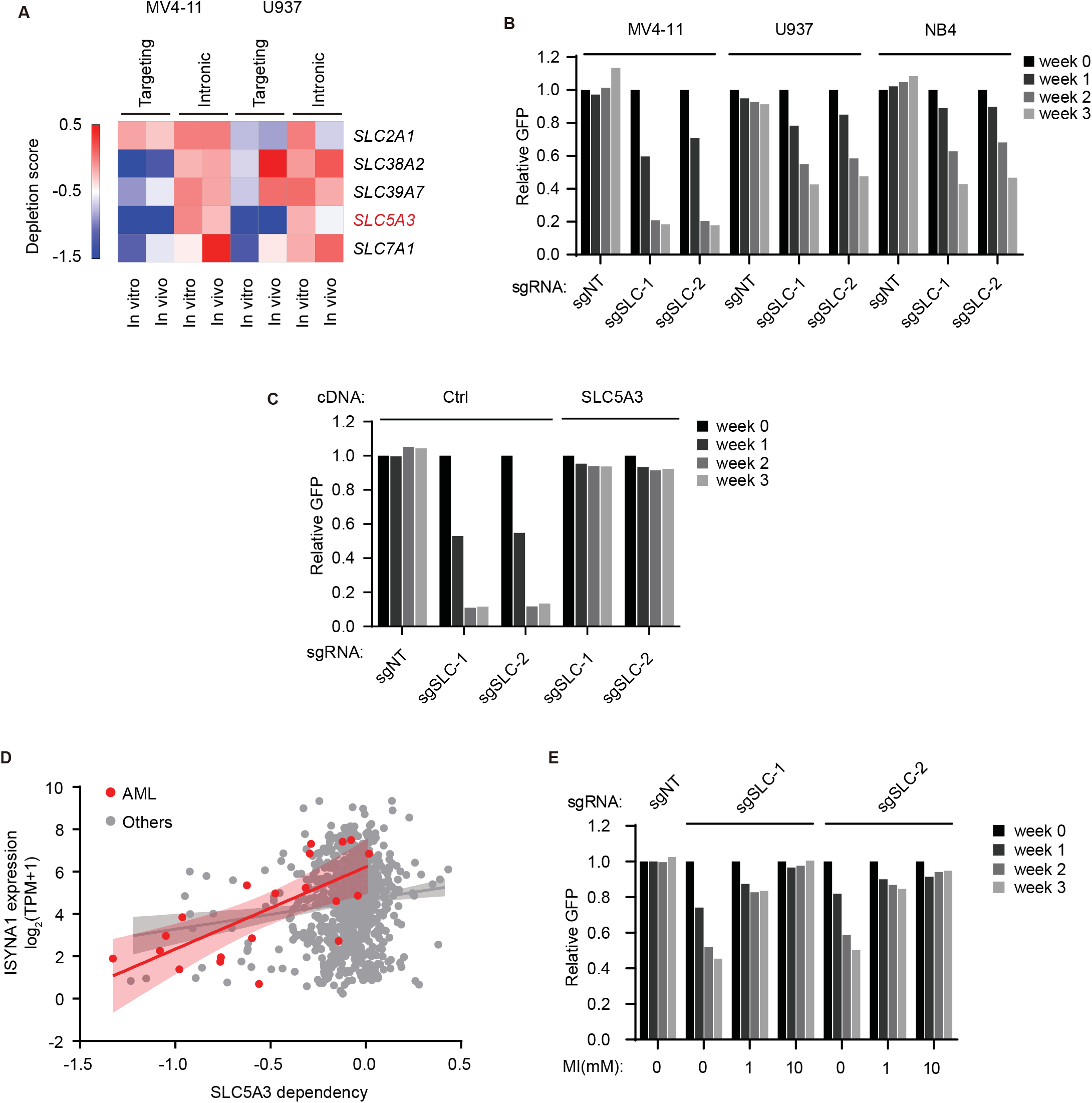
SLC5A3 supports AML growth via myo-inositol transport. **A.** Heatmap showing depletion scores of all solute carrier family genes included in the library. Both targeting and intronic control scores are displayed. **B.** AML cell lines were transduced with non-targeting sgRNA (sgNT) and *SLC5A3* sgRNA (sgSLC-1 and sgSLC-2) vectors that co-express GFP. Cell growth was evaluated in an in vitro competition proliferation assay as measured by the change of percentage of GFP+. **C.** Competitive growth of MV4-11 cells transduced with empty vector (Ctrl) or CRISPR-resistant *SLC5A3* upon endogenous *SLC5A3* knockout. **D.** Scatterplot showing the linear correlation between CERES dependency scores of *SLC5A3* and the expression level of *ISYNA1* across AML cell lines (r=0.700) or other cancer cell lines (r=0.127) in the DepMap dataset. Each dot represents a cell line; the shaded area represents the 95% confidence level interval for the linear model. **E.** Competitive growth was evaluated for *SLC5A3*-knockout MV4-11 cells in culture medium supplemented with indicated concentration of myoinositol (MI). One representative experiment of 2 replicates is shown **B-C** and **E**.

To ensure sufficient library representation in vivo, we first optimized the screening conditions through barcoding experiments using a library of 3152 barcodes. Five to ten million barcoded MV4-11 cells were injected via tail vein into immunodeficient NSGS (NOD scid gamma SGM3) mice. Sub-lethal irradiation was necessary for improved barcode representation and reduced mouse-to-mouse variation (**Figure S2A-B**). Although the barcode distribution was skewed in individual mice even with irradiation, a complete and balanced library representation could be recovered by combining readouts from multiple mice (**Figure S2C-D**). We then performed parallel in vivo and in vitro screens with these optimized conditions using both MV4-11 and U937 xenograft models (**Figure 1B**). Cells were transduced with the screen library in duplicate and selected for two days with puromycin, and then an aliquot of cells was collected as the input reference. We then injected 10 million cells per mouse by tail vein into 4 irradiated mice per replicate and in parallel cultured an aliquot of cells from each replicate in vitro. In vitro cultures were harvested two (for MV4-11) or three (for U937) weeks later, and the bone marrow and spleens were collected when the mice displayed evidence of overt disease. There was strong replicate reproducibility for both the in vitro and in vivo results, although the data generated from U937 displayed a smaller dynamic range likely due to weaker Cas9 activity (**Figure S3A** and data not shown). The average relative abundance of each sgRNA in the output compared to the input samples was determined. The correlation of sgRNA depletion in the bone marrow versus the spleen was strong (**Figure 1C**); therefore, we focused on the bone marrow data for the downstream analysis.

We calculated a normalized depletion score for each sgRNA (see methods, **Supplementary Table 1**). The median value of each set of three sgRNAs was used to represent the score of the corresponding gene. Using the intronic guide population as a null distribution, we defined hits for each cell line. In vitro and in vivo hits were generally well correlated. However, a modest number of targets did not score well in vivo, with a few targets displaying an obvious in vitro versus in vivo discrepancy (**Figure S3B**), underscoring the importance of an in vivo validation strategy for hits emerging from a primary in vitro screen. Supporting the validity of our screen, the genes involved in MLL (mixed lineage leukemia gene, also called *KMT2A)* complex and *FLT3* specifically scored in MV4-11, as expected, since MV4-11, but not U937, is driven by an MLL-fusion oncogene and a FLT3-ITD mutation (**Figure S4A-B**). In addition, several hematopoietic lineage-related transcription factors were strong in vivo dependencies in both cell lines (**Figure S4C**), corroborating recent studies targeting transcriptional vulnerabilities in AML(17,18), and Gene Ontology analysis showed an enrichment of metabolism and mitochondria associated pathways (**Figure S4A and S4D**), consistent with recent findings that AML cells rely on unique metabolic and mitochondrial properties for survival (19,20). Accordingly, our screen provided an informative list of AML targets with high physiological relevance (**Supplementary Table 2**).

Next, we focused on targets that previously had not been described as AML dependencies and ranked highly as in vivo hits in both cell lines: the sodium/myo-inositol cotransporter *SLC5A3* and the mitochondria-localized RING-type ubiquitin E3 ligase *MARCH5* (**Figure 1D-E and Figure S5A-B**)(21,22). We next re-mined the latest edition of the DepMap CRISPR screening datasets, which continue to be expanded, and found that *SLC5A3* and *MARCH5* were indeed strong dependencies in the majority of AML cell lines, displaying a more essential role in AML compared with other cancer types (**Figure S5C**). We therefore selected these two targets for further validation as novel therapeutic opportunities for AML.

### SLC5A3 transports myo-inositol to support AML cell proliferation

*SLC5A3* belongs to the solute carrier family, and among all 5 solute carrier family members included in our screen library, SLC5A3 was the top scoring in both cell lines (**Figure 2A**). We validated this dependency in several AML cells lines using two independent *SLC5A3*-targeting sgRNAs. *SLC5A3* depletion suppressed the growth of AML cells as demonstrated by an in vitro competition assay (**Figure 2B**). The on-target effect was confirmed by using a CRISPR-resistant *SLC5A3* cDNA to rescue the growth defect (**Figure 2C**). Myo-inositol and its derivatives are involved in several cellular processes. While cells take up myo-inositol from the extracellular fluid via both passive and active transportation, they can also synthesize myo-inositol de novo from glucose 6-phosphate. *Inositol-3-phosphate synthase 1 (ISYNA1)* encodes the rate-limiting enzyme in the myo-inositol biosynthesis pathway(23). Intriguingly, the low expression of *ISYNA1* predicted a strong *SLC5A3* dependency in AML cell lines (**Figure 2D**). This correlation suggests that SLC5A3 becomes essential in AML cells with insufficient myo-inositol biosynthesis capacity, and the growth defect caused by SLC5A3 inactivation results from the myo-inositol deficiency. Indeed, with the addition of supplementary myo-inositol in the culture medium, the proliferation of *SLC5A3*-knockout cells was completely rescued (**Figure 2E**). Together, these data reveal that myo-inositol is a critical metabolite for AML, and *SLC5A3* is required for maintaining myo-inositol levels to support AML proliferation.

### MARCH5 loss represses AML cell growth in vitro and in vivo

We next sought to validate the dependency of AML cells on *MARCH5.* Inactivating MARCH5 via either doxycycline (Dox)-inducible CRISPR or shRNA systems induced a severe growth defect in AML cell lines with various genetic backgrounds (**Figure 3A and Figure S6A-B**) (24). The growth defect could be reversed by a CRISPR-resistant cDNA encoding wild-type *MARCH5,* proving the on-target effect. By contrast, *MARCH5* mutations (H43W and C68S) that disrupt its RING domain and thus ubiquitinase function ablated the rescuing ability (25,26), indicating the requirement for the catalytic function of MARCH5 in AML (**Figure 3B and Figure S6C**). Additionally, for MARCH5 validation we deployed a dTAG system, which uses a hetero-bifunctional small molecule that binds the FKBP12^F36V^-fused target protein (i.e., MARCH5) and an E3 ligase complex (i.e., VHL), bringing the two in close proximity and leading to the ubiquitination and proteasome-mediated degradation of the target protein (**Figure S6D**)(27). In this case, we deleted endogenous *MARCH5* by CRISPR and expressed exogenous *MARCH5* with a FKBP12^F36V^-tag, showing that the defective growth can be recapitulated by MARCH5 degradation with the dTAG molecule dTAG^V^-1 (**Figure 3C-D**).

**Figure 3.**
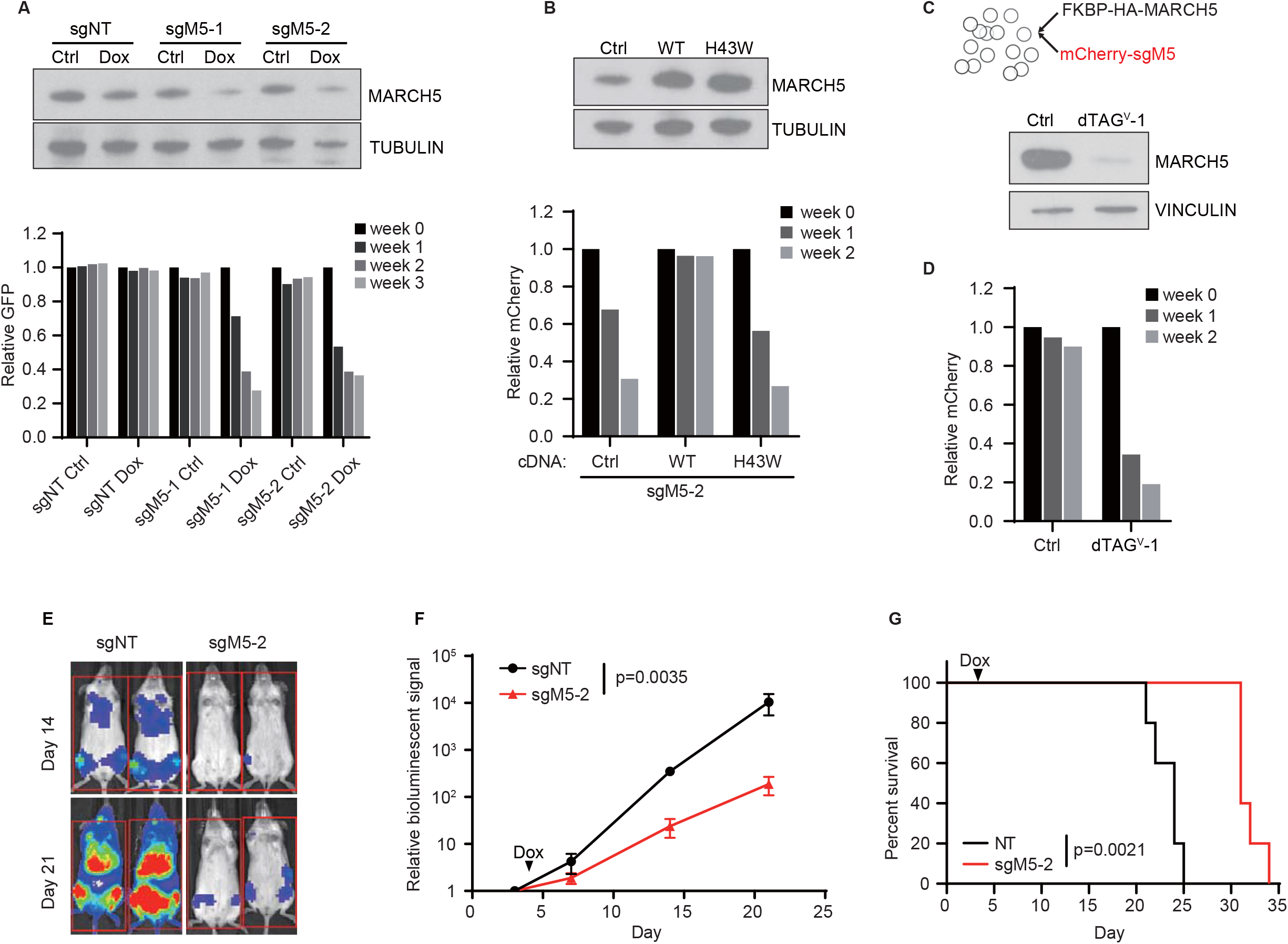
MARCH5 inhibition suppresses AML cell growth. **A.** MV4-11 cells were transduced with Dox-inducible non-targeting sgRNA (sgNT) and *MARCH5* sgRNA (sgM5-1 and sgM5-2) vectors that co-express GFP. Immunoblot analysis of MARCH5 was performed on day 6 after Dox treatment (upper). Cell growth was evaluated in a competition proliferation assay (lower). **B.** Immunoblot analysis of MV4-11 cells expressing an empty vector (Ctrl), *MARCH5* wildtype (WT) or ligase defective mutant (H43W) cDNA (upper). Competitive growth of these cells were evaluated upon endogenous *MARCH5* knockout (lower). **C.** Schematic of the MARCH5 dTAG system (upper). Immunoblot analysis of FKBP12^F36V^-tagged MARCH5 with HA antibody in NB4 cells treated with 500 nM dTAG^V^-1 for 24 hours (lower). **D.** Competitive growth of NB4 MARCH5-dTAG cells treated with DMSO (Ctrl) or 500 nM dTAG^V^-1. **E.** NSGS mice were transplanted with MV4-11 cells expressing Dox-inducible sgNT or *MARCH5* sg-2. Dox-containing food was served from day 4 post transplantation. Representative bioluminescence imaging images are shown on the indicated day post transplantation. **F.** Quantification of serial bioluminescence imaging. The data were normalized to the baseline readout on day 3. n=5, results represent mean ± SD. The p-value was calculated using unpaired two-tailed t-test with measurements on day 21. **G,** Survival curves of mice used in (**F**). The p-value was calculated by log-rank test. One representative experiment of 3 replicates is shown in **A**, **C** and **D**, of 2 replicates is shown in **B**.

Using the luciferase-expressing MV4-11 cells, we confirmed that Dox-mediated deletion of *MARCH5* post transplantation led to a marked attenuation of AML progression in NSGS mice as monitored by bioluminescence imaging, which translated to prolonged survival (**Figure 3E-G**). Importantly, these results also demonstrate that the in vivo growth disadvantage of *MARCH5*-depleted cells is not caused by homing defects but rather that MARCH5 is critical for disease progression *in vivo.*

### MARCH5 prevents apoptosis in AML

We observed that MARCH5 inactivation led to apoptosis induction in AML cells as indicated by upregulated cleaved-Caspase3 and Annexin V (**Figure 4A-B and Figure S7A**). Importantly, knockout of the mitochondrial apoptosis effectors *BAX* or *BAK1* reversed the apoptosis induction and growth defect of *MARCH5*-null cells (**Figure 4C-D and Figure S7B-C**). Interestingly, AML cell lines displayed differential reliance on *BAX* and *BAK1* for the execution of MARCH5-depletion mediated apoptosis. In contrast to MV4-11 and MOLM14 cells, double-knockout of *BAX* and *BAK1* was required to rescue MARCH5 inactivation in NB4 cells (**Figure 4E-F and Figure S7D**). Nevertheless, these results demonstrate that apoptosis induction is an essential cellular mechanism accounting for the inhibitory effect of MARCH5 ablation in AML.

**Figure 4.**
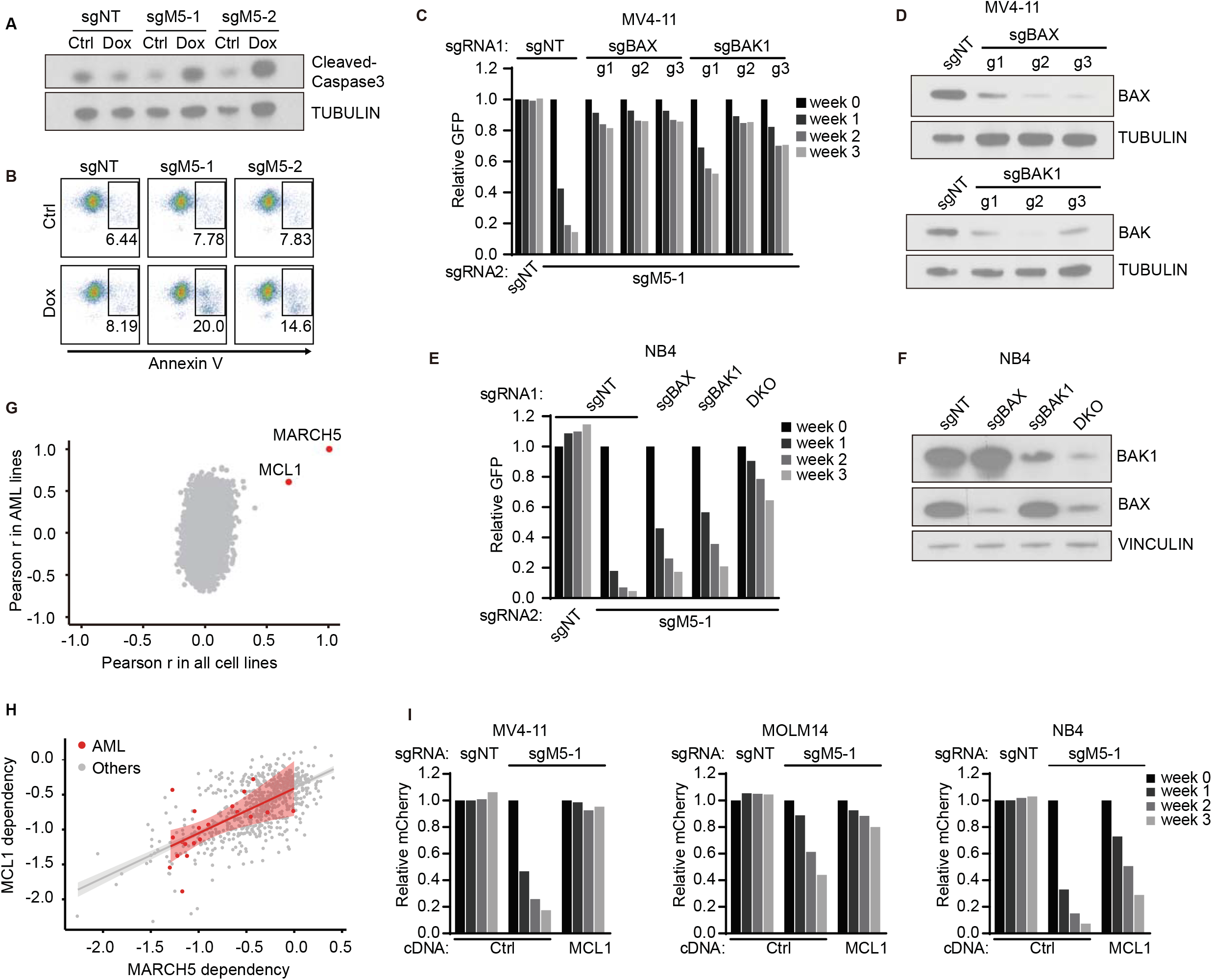
Inhibition of MARCH5 activates mitochondrial apoptosis pathway. **A, B.** Immunoblot analysis of cleaved-caspase3 (**A**) and flow cytometry analysis of Annexin V (**B**) in MV4-11 cells transduced with the indicated sgRNAs at day 10 after Dox treatment. **C.** Competitive growth was evaluated for the control or *BAX/BAK1-* knockout MV4-11 cells, which were generated with 3 independent sgRNAs each, upon *MARCH5* deletion. **D.** Immunoblot analysis of BAX/BAK1 in MV4-11 cells described in (**C**). **E,** Competitive growth assay for control, *BAX*-, *BAK1-* and double (DKO)-knockout NB4 cells with *MARCH5* depletion. **F.** Immunoblot analysis of BAX/BAK1 in NB4 cells described in (**E**). **G.** Scatterplot showing the Pearson correlations between CERES dependency scores of *MARCH5* and each other gene across AML cell lines or all cancer cell lines in DepMap CRISPR screen dataset. Each dot represents a gene. **H.** Scatterplot showing the linear correlation between CERES dependency scores of *MARCH5* and *MCL1* across AML cell lines (r=0.609) or other cancer cell lines (r=0.657) in the DepMap dataset. Each dot represents a cell line; the shaded area represents the 95% confidence level interval for the linear model. **I.** Competitive growth of control or *MCL1*-overexpressing cells with *MARCH5* knockout (sgM5). One representative experiment of 2 replicates is shown in **A-F** and **I**.

The mitochondrial apoptotic pathway is regulated by interactions among BCL2 family proteins(28). Given the association of MARCH5 depletion with apoptosis in AML, we explored the possible crosstalk between MARCH5 and BCL2 family proteins. Strikingly, multiple genome-wide screens revealed that the dependency scores of *MARCH5* and *MCL1*, but not other anti-apoptotic genes, were significantly correlated across the AML cell lines as well as other cancer lines (**Figure 4G-H and Figure S7E**), suggesting that *MARCH5* and *MCL1* are functionally connected. We hypothesized that enhancing the MCL1 activity can reverse the effect of MARCH5 inhibition. Indeed, overexpression of *MCL1* completely or partially rescued the growth impairment observed with MARCH5 depletion in AML cells (**Figure 4I**), supporting the notion that loss of MARCH5 may disrupt the function of MCL1.

### MARCH5 depletion sensitizes AML cells to venetoclax

Apoptosis sensitivity can determine the response to various therapies in cancer (29). Because loss of MARCH5 primed AML cells to apoptosis, we investigated whether MARCH5 inhibition can sensitize AML cells to anti-cancer therapies. Control and MARCH5-knockdown OCI-AML2 cells were subjected to a chemical screen across a library of 3247 anti-cancer compounds. Fifty-eight compounds displayed an enhanced inhibitory effect on MARCH5-knockdown cells compared to control cells. Notably, the BH3-mimetics, a class of small molecules that mimic BH3-proteins to bind and inhibit anti-apoptotic BCL-2 family proteins, were enriched (**Figure 5A-C and Figure S7F**). These data prompted us to examine whether MARCH5 inactivation in AML cells can enhance their sensitivity to venetoclax, a BH3-mimetic that specifically blocks BCL2 and is FDA-approved for the treatment of older adults with AML in combination with a hypomethylating agent (30,31). MARCH5 depletion through inducible CRISPR or dTAG degradation indeed sensitized AML cells to venetoclax (**Figure 5D-E**). The cooperativity between MARCH5 inhibition and venetoclax was also demonstrated in the xenograft model of luciferase-expressing MV4-11 cells. Although the venetoclax regimen used showed minimal impact on control cells, it enhanced the anti-leukemic activity of MARCH5-depletion in vivo (**Figure 5F-G**). All told, these results highlight the combinational targeting of MARCH5 and BCL2 as a potential therapeutic approach for AML.

**Figure 5.**
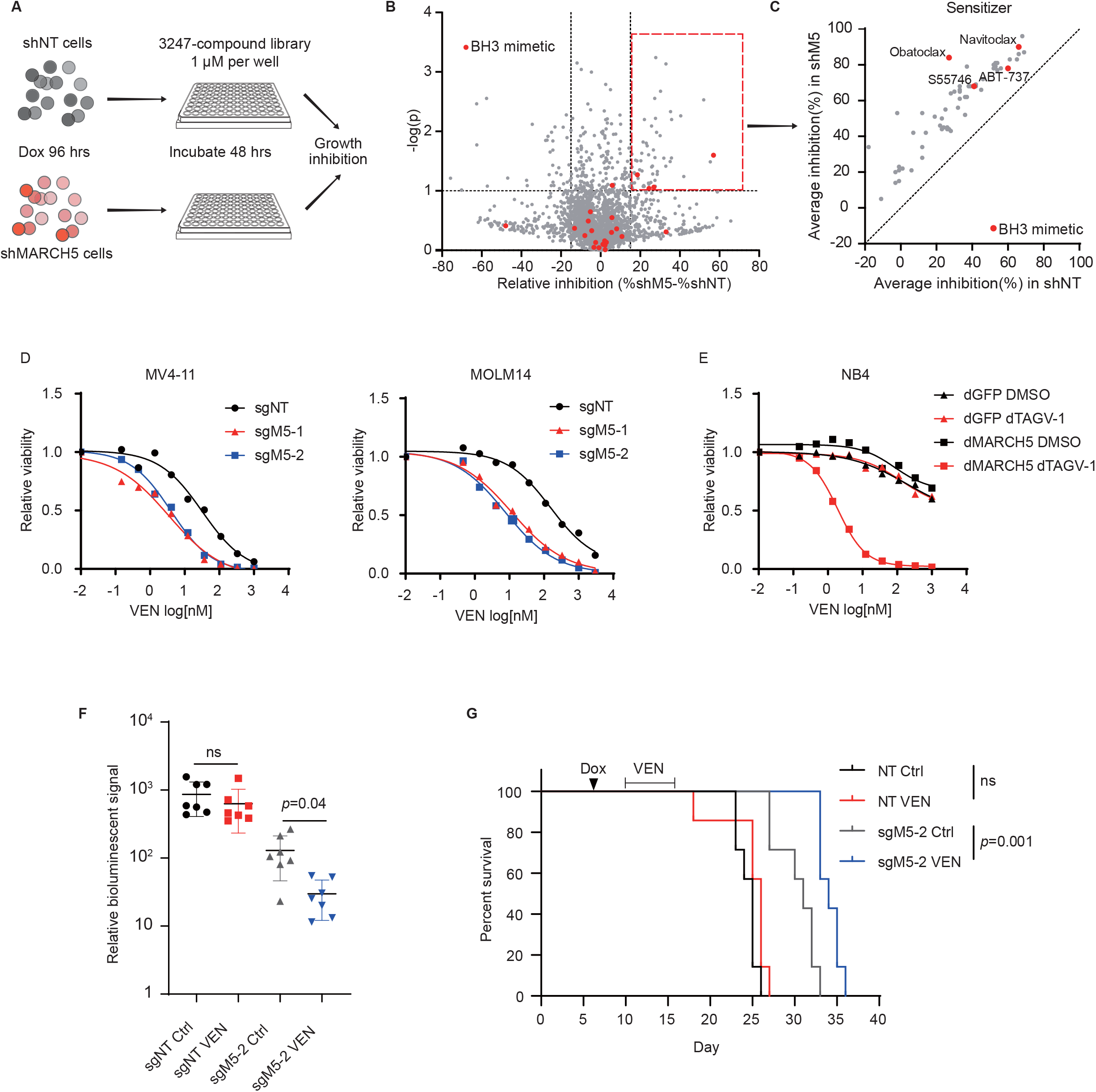
Inactivating MARCH5 enhances the anti-leukemic activity of venetoclax. **A.** Schematic of chemical screen, 1 μM of each compound was used in the screen. **B, C.** Scatterplot showing the relative inhibitory effect of screening chemical compounds in MARCH5-knockdown cells (shM5) compared to control cells (shNT) (**B**). Each dot represents a compound. The relative inhibition is the mean difference of percentage of growth inhibition between shM5 and shNT cells. *p* values were calculated by unpaired two-tailed t-test, n=2. A cutoff of *p*≤0.1 and relative inhibition?l5% was used to select compounds displaying the enhanced inhibitory effect on shM5 cells, which were present in (**C**). Compounds within the class of BH3-mimetics are highlighted. **D, E.** Relative proliferation of Dox-inducible sgRNAs-(**D**) or FKBP-MARCH5 (dM5)-expressing cells (**E**) treated with venetoclax (VEN) for 3 days. Cells were treated with Dox for 4 days prior, or 500 nM dTAG^V^-1 concurrently with venetoclax treatment. Cells expressing FKBP-GFP (dGFP) were included as a control. Cell viability was determined by CellTiter-Glo, and normalized to the DMSO-treated control. The mean (n=8) and four-parameter doseresponse curves are plotted. **F.** NSGS mice were transplanted with MV4-11 cells expressing Dox-inducible sgNT or *MARCH5* sg-2. Dox-containing food was served from day 7 post transplantation. Mice were treated for one week with 75 mg/kg of venetoclax by oral gavage daily starting day 10 post transplantation. Quantification of bioluminescence imaging on day 18 is shown. The data were normalized to the baseline readout on day 3. n=7, results represent mean ± SD. The p-values were calculated using unpaired two-tailed t-test. **G.** Survival curves of mice used in (**F**). The p-values were calculated by log-rank test. For **D-E**, one representative experiment of 2 replicates is shown.

### Dependency validation in CRISPR-competent PDX models

To further evaluate the therapeutic potential of the identified AML dependencies, we investigated their essentiality in PDX models, argued to be the most faithful models to primary human disease(11). CRISPR-mediated genetic studies in PDXs have been challenging due to the poor transduction efficiency of PDX cells and limited growth in vitro. To develop PDX models that are tractable for CRISPR-editing, we screened a cohort of PDX samples and identified those transducible and suitable for short-term in vitro culture (**Supplementary Table 3**). These PDX cells were transduced with lentivirus co-expressing Cas9 and a fluorescent protein (GFP or mCherry). Cas9-expressing PDX cells were purified based on fluorescence and expanded via serial transplantation into NSGS mice (**Figure 6A-B**). Cas9 activity was assessed using a fluorescent protein-linked sgRNA targeting *CD33.* More than 80% of PDX cells receiving the sgRNA became CD33 negative, indicating a high Cas9 activity in these models (**Figure S8A**). Importantly, CRISPR-knockout of *SLC5A3* and *MARCH5* dramatically reduced in vitro cell growth of PDX16-01(complex karyotype, *TP53* and *PTPN11* mutations) and PDX17-14 (complex karyotype and MLL-AF10), highlighting their critical roles in AML (**Figure 6C-D**).

**Figure 6.**
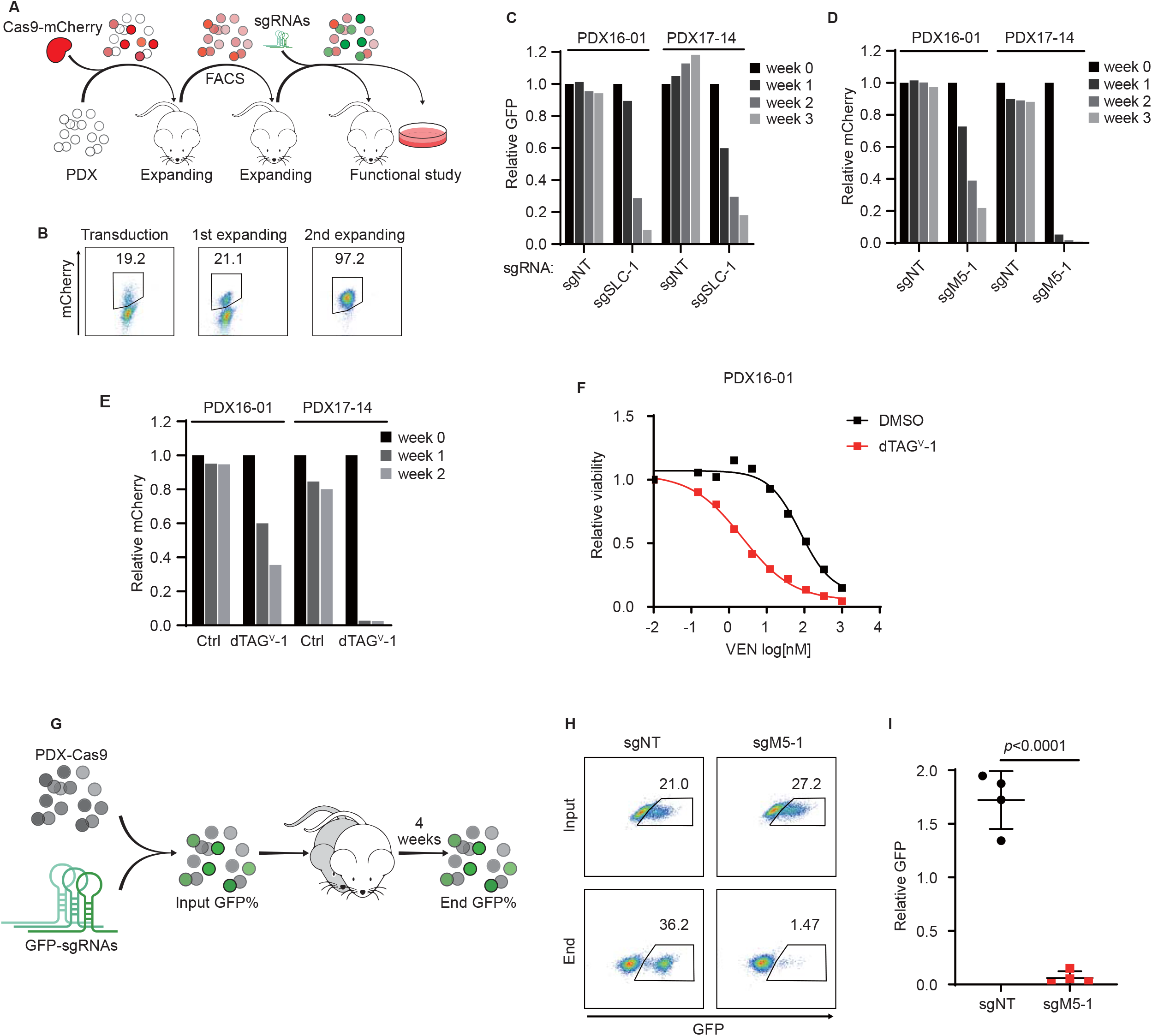
Dependency validation in AML PDX models. **A.** Schematic for generating PDX models capable of CRISPR-editing. PDXs expressing mCherry or GFP-linked Cas9 were established. **B.** Representative flow cytometry analysis showing the purification and expansion of Cas9-mCherry-expressing PDX cells. **C, D** Competitive growth of Cas9 PDX cells transduced with fluorescent protein-linked sgNT or sgRNA targeting *SLC5A3* (sgSLC-1 in **C**) or MARCH5 (sgM5-1 in **D**). **E.** Competitive growth of MARCH5-dTAG PDX cells treated with DMSO (Ctrl) or 500 nM dTAG^V^-1. **F.** Relative proliferation of MARCH5-dTAG PDX cells treated with venetoclax for 3 days. The mean (n=8) and four-parameter dose-response curves are plotted. **G.** Schematic of in vivo competition assay with PDX cells. Mouse bone marrow cells were collected for evaluating end GFP+ percentage. **H.** Representative flow cytometry analysis of input and end GFP percentage of sgNT-or *MARCH5* sg-1-expressing cells. **I.** Relative abundance of PDX cells expressing sgNT or *MARCH5* sg-1, as calculated by normalizing end GFP percentage to input GFP percentage. The p value was calculated by unpaired two-tailed t-test, n=4. For **B-F**, one representative experiment of 2 replicates is shown.

We focused on MARCH5, and established a MARCH5 dTAG degradation system in PDX models (**Figure 6E**). In PDX17-14, which is highly sensitive to MARCH5 inhibition, 2-hour treatment with dTAG^V^-1 was sufficient to prime cells for apoptosis as indicated by BH3-profiling (**Figure S8B**)(29). Consistent with the observation in cell lines, the apoptosis induced in PDX cells can be rescued by deletion of *BAX* and/or *BAK1* (**Figure S8C**), and PDX cells without MARCH5 became more sensitive to venetoclax (**Figure 6F**). Importantly, the MARCH5 dependency was further confirmed in an in vivo competition assay using a third PDX model (PDX68555 with complex karyotype and MLL-AF9). PDX cells expressing a GFP-linked sgRNA targeting *MARCH5* were depleted in NSGS mice, as evidenced by a dramatic reduction of the GFP+ fraction in engrafted cells. In contrast, the PDX cells expressing a non-targeting sgRNA were maintained (**Figure 6G-I**). These results demonstrate that targeting MARCH5 can suppress the progression of AML PDX cells both in vitro and in vivo. Collectively, our CRISPR-competent PDX models provide a unique opportunity to study genetic dependencies of AML within a clinically relevant context.

## Discussion

In vivo CRISPR screens offer a strategy for identifying novel therapeutic targets for leukemia within the context of the physiologic microenvironment. Previous studies have adapted this approach in a mouse MLL-AF9-driven AML model and a mouse BCR-ABL/NUP98-HOXA9-driven chronic myeloid leukemia model to identify genes essential for tumor growth(8,9). Here, we developed an in vivo CRISPR screening pipeline in orthotopic xenograft models of human AML, providing a complementary approach for AML dependency identification. We have defined experimental conditions necessary for an optimal in vivo screen. One major consideration is the requirement for sufficient in vivo library representation to avoid false positive hits. Since AML is established by leukemia initiating cells, only a subset of the AML cell population retains this leukemia initiating activity in mouse recipients. As evidenced by our barcoding experiments, random sets of barcodes prevailed in different individual recipients, reflecting the selective leukemia initiating activity and clonal expansion. Therefore, it is critical to utilize irradiation to maximize the engraftment capacity of AML cells and include multiple mice to achieve complete in vivo library representation. The number of mouse recipients has to be adjusted according to the library size and the leukemia initiating capacity of the model.

Large-scale genome-wide CRISPR screening has provided numerous potential AML targets. Our study aimed to refine this candidate list by providing in vivo functional references, which will inform future studies. We observed a strong correlation between the in vitro and in vivo dependencies, likely related to the reduced off-target effects of CRISPR compared to other approaches such as RNAi and the robust library representation. However, a few genes reproducibly appeared as in vitro-only dependencies, demonstrating that the microenvironment can influence the essentiality of a target. Due to the design of the screening library, prioritized based on in vitro hits, our current study was not positioned to discover in vivo-specific dependencies. Coupled with other focused libraries, however, our pipeline is adaptable for de novo dependency discovery in human AML.

Metabolic reprogramming contributes to tumor development and sustains cancer cell proliferation. Like other cancers, AML has altered metabolic features and is addicted to certain metabolites and metabolic pathways for survival. For instance, the vitamin B6 pathway is reported to be critical for AML growth(32), and AML stem cells selectively depend on amino acid metabolism to fuel oxidative phosphorylation(33). Such metabolic addictions also provide new possibilities for AML treatment. For example, blocking the one-carbon folate and oxidative phosphorylation pathways has exhibited anti-leukemic activity in preclinical studies(33,34). Here, we revealed myo-inositol as a metabolic dependency in AML. Our data suggest that AML cells rely on either SLC5A3-mediated extracellular transportation or ISYNA1-mediated intracellular de novo synthesis for their myo-inositol supply. AML cells with low ISYNA1 activity depend on SLC5A3 exclusively for fueling myo-inositol metabolism. Interestingly, the SLC5A3-ISYNA1 correlation is less evident in other cancer types; the majority of ISYNA1 low-expressing cancer cell lines are not dependent on SLC5A3. The underlying biology rendering SLC5A3 as a selective dependency in AML with low ISYNA1, but not in all cancers with low ISYNA1, is not clear. It is possible that the myo-inositol addiction is endowed by the cell lineage in AML or that alternative myo-inositol transporters are utilized in ISYNA1 low-expressing cancer cells that are not SLC5A3 dependent. Although SLC5A3-specific inhibitors have not been reported, small molecule inhibitors against SLC5A1 and SLC5A2, two solute carrier family 5 members with a similar structure to SLC5A3, have been developed(35,36). Therefore, SLC5A3 is potentially actionable, and our study supports the development of chemical probes against this strong AML dependency.

Mitochondrial metabolism is another important metabolic axis in AML disease maintenance and drug response/resistance. It is therefore particularly relevant that MARCH5, a target located in the outer mitochondrial membrane, scored as a top hit in our screens. MARCH5 has been reported to serve multiple context-dependent functions in cells, including regulation of mitochondrial dynamics through its modulation of fission effector proteins such as DRP1 and MID49, or fusion effector proteins such as mitofusin 1 and mitofusin 2 (MFN1 and MFN2) (22); protection against stress stimuli, including endoplasmic reticulum stress, antimitotic drugs, and BH3-mimetics through facilitation of mitochondria hemostasis or regulation of stress-responding proteins such as inositol-requiring kinase 1 (IRE1) and NOXA (26,37–40); and prevention of persistent innate immune response through the reduction of mitochondrial antiviral signaling (MAVS) aggregates(41). In contrast, in the case of AML, we have revealed that MARCH5 is essential for cell survival under physiological conditions; inhibiting MARCH5 by itself is sufficient to activate the canonical mitochondrial apoptosis pathway in a BAX/BAK1-dependent manner. The dependency correlation analysis and our functional studies strongly suggest that MARCH5 regulates apoptosis via interfering with MCL1 function. While we cannot rule out a potential role for altered mitochondrial dynamics in the AML phenotype observed with MARCH5 depletion, dependency scores for mitochondria dynamics genes were not significantly correlated with a MARCH5 dependency in the DepMap dataset.

Venetoclax has received FDA approval for the treatment of newly diagnosed older adult patients with AML in combination with hypomethylating agents. This promising combination has resulted in a remission rate of approximately 70%. However, intrinsic and acquired resistance still emerged in a significant percentage of patients, and the efficacy of this combination drops precipitously in patients with relapsed and refractory AML(30,31). Thus, additional combination approaches are desired to enhance the clinical benefits of venetoclax. Overcoming venetoclax resistance via targeting mitochondrial components is an emerging theme. Blockage of oxidative phosphorylation activity, inhibition of mitochondrial translation, and disruption of mitochondrial cristae structures can all result in venetoclax sensitization in AML cells(33,42,43). Consistent with previous reports showing loss of MARCH5 sensitizing cells to BH3-mimetics(26,38), our data have demonstrated that MARCH5 is another promising synergistic mitochondrial target for enhancing the efficacy of venetoclax in AML. Upregulation of MCL1 activity has been identified as a mechanism causing venetoclax resistance(44), consistent with MARCH5 depletion mediating venetoclax response by impairing MCL1 function. Notably, in addition to BCL2 family inhibitors, MEK inhibitors were enriched in our MARCH5 depletion chemical sensitizer screen. It has been reported that MEK1/2 inhibitors are synthetic lethal with MCL1 inhibitors in treating melanoma(45). Similar crosstalk may also occur here, again suggesting an impaired MCL1 activity in MARCH5-depleted AML cells. Collectively, MARCH5 is positioned as a strong AML dependency, as well as a synergistic target for anti-BCL2 therapy. With the success of targeting other E3 ligases (e.g., MDM2 and XIAP) by small molecules already in clinical testing in humans(46,47), and given that the enzymatic activity of MARCH5 is critical, MARCH5 targeting holds promise for patients with AML and potentially the treatment of other malignancies.

PDXs provide useful pre-clinical models for therapy evaluation. These models largely retain the histologic and genetic characteristics of the primary disease and have also been shown to be predictive of clinical outcome(11). While PDX models have been widely used for drug evaluation, we established the CRISPR-competent PDX models to enable functional genetic studies. In addition, we demonstrated the utility of dTAG-directed protein degradation for mimicking pharmacological inhibition of the target. These tools are a valuable addition to the AML dependency prioritization and validation, especially for studying targets without a tool compound inhibitor available, such as SLC5A3 and MARCH5. Future studies will investigate whether the dependency landscape obtained from the cell lines is broadly reflected in PDXs.

Taken together, our in vivo screening approach, coupled with CRISPR-competent PDX models and dTAG-based degrader systems, constitute a platform for prioritizing and identifying AML targets for therapy development, with SLC5A3 and MARCH5 nominated as two top targets for further consideration in AML.

## Methods

### Plasmids and reagents

The lentiviral Cas9 vectors co-expressing a blasticidin resistance gene or GFP (pXPR_BRD101 and pXPR_BRD104) and the lentiviral sgRNA expression vectors coexpressing a puromycin resistance gene, mCherry or mAmetrine (pXPR_BRD003, pXPR_BRD043 and pXPR_BRD052) were provided by the Genetic Perturbation Platform (GPP) at the Broad Institute. The Cas9-T2A-mCherry and Dox-inducible sgRNA vectors were obtained from Addgene (#70182 and #70183). sgRNAs were cloned into BsmBI-digested vectors. shRNA sequences were constructed into pLKO-TET-ON (Addgene # 21915). Sequences of sgRNA and shRNA are provided in **Supplementary Table 4**. For overexpression, MARCH5 and MCL1 cDNAs were synthesized as gBlocks fragments (Integrated DNA Technologies), and then cloned into the lentiviral expression vectors coexpressing a puromycin resistance gene or GFP (pLX_TRC307 and pLX_TRC312 from GPP) using a Gibson Assembly Cloning Kit (New England Biolabs E5510S). dTAG expression vector and dTAG^V^-1 compound were kindly provided by Dr. Nathanael Gray. Venetoclax was acquired from Selleck (S8048) or MedChemExpress (HY-15531).

### Cell culture and patient-derived xenograft samples

AML cell lines (MV4-11, U937, MOLM14, NB4 and P31FUJ) were cultured in RPMI supplemented with 10% fetal bovine serum (FBS) and 1% penicillin-streptomycin (PS). HEK293T cells were cultured in DMEM with 10% FBS and 1% PS. All cell lines have been STR-profiled at the Dana-Farber Cancer Institute.

Primary patient samples were acquired following informed consent and patient-derived xenografts (PDXs) were established under protocols approved by Dana-Farber Cancer Institute and Cincinnati Children’s Hospital Medical Center institutional review boards. The detailed information on PDXs is provided in **Supplementary Table 3**. For short-term in vitro culture, PDX cells were maintained in IMDM containing 20% FBS and 1 % PS, and supplemented with 10 ng/mL human SCF, TPO, FLT3L, IL-3 and IL-6 (PeproTech 300-07, 300-18, 300-19, 200-03 and 200-06).

### Lentivirus production and transduction

Virus was produced using HEK293T cells transfected with lentiviral expression vectors, together with envelope VSVG and the gag-pol psPAX2 constructs. For transduction, AML cells were mixed with viral supernatant and 4-8 μg/mL polybrene. In some experiments, cells were centrifuged in viral supernatant at 1000 g for 1hr at 33 °C to enhance the transduction efficiency.

### Flow cytometry analysis

Apoptosis was analyzed with an APC Annexin V Apoptosis Detection Kit (BioLegend) following the manufacturer’s protocol. Cells from mouse tissues were incubated with nonspecific binding blocker (anti-mouse/human CD16/CD32 Fcγ receptor, BD Biosciences) before staining for APC-human CD33 (Biolegend #303408), V450-human CD45 and APC-Cy7 mouse CD45 (BD Biosciences #560368 and #557659). Cells were analyzed on an LSRFortessa or FACSCelesta flow cytometer or sorted on a FACSAria II flow cytometer (BD Biosciences) or a SH800S flow cytometer (Sony), and the data was analyzed with FloJo software (TreeStar).

### Competition assay

Cells were transduced with lentivirus vectors co-expressing a sgRNA and a fluorescent protein, such as GFP, mCherry or mAmetrine, at an efficiency of approximately 50%, or the transduced cells were mixed with non-transduced cells at approximately a 1:1 ratio. The cell growth was evaluated by the change in the fraction of cells expressing the fluorescent protein, which was monitored by flow cytometry. For the dTAG experiment, 500 nM dTAG^V^-1 was added into culture and replenished every 3-4 days.

### Venetoclax treatment

Cells were plated in 384-well plates at 1,000-1,500 cells per well and mixed with serially-diluted concentrations of venetoclax or 0.1% DMSO as a control. The viability of cells was measured after 3 days incubation using a CellTiter-Glo Luminescent Cell Viability Assay kit (Promega) following the manufacturer’s protocol. Data were analyzed using GraphPad Prism software.

### Xenograft transplantation

Transplantation was performed on 6-to 8-week-old NOD/SCID/IL2RG-/- immunodeficient mice with transgenic expression of human SCF, GM-CSF and IL-3 (NSGS, Jackson Laboratory). When necessary, mice were conditioned with sublethally irradiation at least 6 hours before transplantation. For Dox-inducible sgRNA experiments, 2×10^5^ luciferase expressing MV4-11 cells were transplanted into each mouse via tail vein. Dox-containing food was initiated on day 4 post transplantation. Leukemia progression was serially assessed using bioluminescence imaging. Mice were injected with 75 mg/kg i.p. d-Luciferin (Promega), anesthetized with 2–3% isoflurane, and imaged on an IVIS Spectrum (Caliper Life Sciences). A standardized region of interest (ROI) encompassing the entire mouse was used to determine total bioluminescence flux. For the experiment evaluating venetoclax treatment in combination with MARCH5 depletion, Dox-containing food was initiated on day 7 post transplantation, and 10 days post injection, mice were treated with vehicle (60% phosal 50 propylene glycol, 30% polyethyleneglycol-400, and 10% ethanol) or venetoclax (75 mg/kg body weight) daily by oral gavage for one week.

For the PDX experiments, 0.5-1×10^6^ AML PDX cells were transplanted. Engrafted cells were collected for analysis when mice displayed overt disease. All experiments were performed in accordance with Dana-Farber Cancer Institute institutional guidelines.

### Western blotting

Whole cell lysate were obtained by directly lysing cells in 2x Laemmli Sample Buffer (Bio-Rad, #1610737) containing reducing reagent and boiling. The lysates were resolved in SDS-PAGE, followed by transfer to nitrocellulose membranes and immunoblotting. The primary antibodies used were anti-MARCH5 (Abcam ab174959, MilliporeSigma 06-1036 and Cell Signaling Technology #19168), anti-TUBULIN (MilliporeSigma T0198) and anti-Vinculin (Abcam ab130007). And anti-HA (#3724), anti-cleaved Caspase3 (#9664), anti-MCL1 (#94296 and #5453), anti-BAX (#5023) and anti-BAK1 (#12105) were all from Cell Signaling Technology. The secondary antibodies were HRP-linked goat anti-rabbit IgG and anti-mouse IgG (Cell Signaling Technology #7074, #7076).

### Generation of AML-enriched dependency library

AML-enriched dependencies were identified in a provisional internal DepMap Avana CRISPR-Cas9 screen dataset consisting of 578 cancer cell lines including 17 AML cell lines (available at https://figshare.com/s/796ac27867ee3f12f2d2). The gene effect scores summarizing the guide depletion for each gene were determined by the CERES algorithm as previously described (4). Probabilities of dependency were calculated for each gene as the probability that the gene effect score arises from the distribution of essential gene scores rather than nonessential gene scores as previously described(48).

Genes were selected based on the following criteria:

1. A two-class comparison of gene effect scores of each gene in AML cell lines (n=17) compared to all other cancer lines (n=561) was performed using the *lmFit* and *eBayes* functions implemented in the *limma* R package. Briefly, *lmFit* was used to fit a linear model to the gene effect scores divided in the in-group and out-group. Then, *eBayes* was used to compute t-statistics and log-odds ratios of differential gene effect. Two-sided p-values were corrected for multiple hypothesis testing using the Benjamini-Hochberg correction and these adjusted p-values were reported as q-values. Genes with significantly lower gene effect scores (greater dependency) in AML (q-value <0.2) compared to other cancer cell lines were identified as candidate AML dependencies for validation.
2. Dependencies were selected that had a false discovery rate (FDR) <0.1 in at least one of these cell lines, MV4-11, MOLM14, U937 and OCIAML3. FDR is calculated as the expected number of false positives by taking the sum of probabilities that a gene is not a dependency (probability of not dependency = 1 – probability of dependency) of all genes with more negative gene effect scores than the gene of interest in a particular cell line. We focused on the 4 cell lines listed above because they have favorable in vivo properties, such as reproducible bone marrow engraftment and rapid disease progression, and they represent a variety of AML genotypes, including with or without a MLL-fusion oncogene.
3. Gene dependencies were also filtered to include AML-enriched dependencies in the DEMETER2 RNAi screen with a q-value <0.3 using the same two-class comparison described above if the gene was present in the RNAi dataset(14).
4. For genetic dependencies that were enriched for depletion in AML compared to other cancer cell lines, we also required that they be depleted in the independent genome-scale CRISPR-Cas9 screen conducted in AML cell lines by Wang et al.(13). The depletion was empirically defined as a log2 fold change (LFC) <-2.26, which is 80 percent of the median LFC of essential genes included in the Wang et al. dataset. The list of essential genes was derived from Hart et al.(3).
5. Genes were then filtered to include those with detectable expression in primary AML samples, with log2 (1+FPKM) > 1 in the TCGA LAML dataset, or log2 (RMA)>6 in GSE14468, or log2 (RMA)>6 in the TARGET AML dataset.
6. Next, we defined common essential dependencies as those have a probability of dependency greater than 0.5 in more than 80% of the 578 cell lines screened and also defined by Hart et al.(3). Common essential genes were filtered out from the list unless they were among the top 40 AML-enriched dependencies ranked by q-value. As such, 21 common essential genes were kept in the library, which were used as the normalization reference for calculating the depletion scores. The selection criteria to this point produced a list of 183 candidate genetic dependencies.
7. Finally, we included MLL-fusion enriched dependencies by comparing 6 MLL-fusion positive AML lines (MOLM13, MONOMAC1, MV411, NOMO1, OCIAML2, THP1) to 8 AML cell lines without MLL-fusion (AML193, F36P, HEL9217, HEL, KASUMI1, NB4, OCIAML3, TF1) using the two-class comparison described above. The 17 most enriched MLL-fusion dependencies that were not already on the gene list described above were added to create a library of 200 potential genetic dependencies for validation.

Three targeting sgRNAs were designed for each gene using the previously described method maximizing Rule Set 2 scores and minimizing off-target sites with high Cutting Frequency Determination scores(49). Additionally, three sgRNAs targeting intronic regions of each gene were generated using the same method. Intronic regions were determined using the shared intronic regions across multiple isoforms using NCBI Refseq UCSC, NCBI RefSeq UCSC, NCBI REfSeq All, GENGODEv24 knownGene. The regions within 30bp to known splice sites or overlapping with an exon of another gene were avoided. To assemble the library, sgRNAs were cloned into the pXPR_BRD003 vector that expresses a puromycin resistance gene as previously described(49).

### In vivo and in vitro screens

Screens were performed in duplicate. MV4-11 or U937 cells stably expressing Cas9 were transduced with the screen library at an efficiency of 30-50%, so that the majority of cells receive only one sgRNA. Puromycin-selection was initiated on the day post transduction for two days, and the selected cells were recovered in fresh medium for one more day. On day 4 post transduction, 1.5 million selected cells, sufficient for a representation of more than 1000 cells per guide, were collected as the input reference. The remaining cells were divided into the in vivo and vitro screens. For the in vivo screen, 10 million cells were transplanted into each sub-lethally irradiated NSGS mouse via tail vein, with 4 mice per replicate. Cells from mouse bone marrow and spleen were collected at around week 3 post the transplantation, when overt disease was observed. For the in vitro screens, at least 1.5 million cells was maintained throughout the 14 to 21-day culture period and collected at the end of the screen. Genomic DNA was extracted from collected cell pellets using the Qiagen DNeasy Blood and Tissue kit (# 69506). The sgRNA barcode was PCR amplified and submitted for standard Illumina sequencing as previously described(4). The barcoding experiments were performed in a similar manner except using a barcoding library.

### Data analysis of CRISPR screens

In each sample, the read counts of each sgRNA were normalized to the total reads per million of all negative control sgRNAs. For in vivo samples, the normalized counts from all 4 mice within a replicate were averaged. The fold-change of each sgRNA in post-screen samples was determined relative to the input for each replicate, which were then log2 transformed and averaged across the two replicates (LFC). The depletion score of each sgRNA was calculated as −1 * (each_guide_LFC-median(negative_controls_LFC)) /(median(common_essentials_LFC)-median(negative_controls_LFC)), which sets the median of negative control targeting guides to 0 and the median of common essential guides to −1. The median value of each 3-sgRNA set was chosen to represent the depletion score of the corresponding genes or intronic controls. The depletion scores of all intronic sgRNAs were used to estimate the null distribution. A target with a depletion score ≤ mean (intronic)-1.28*SD (intronic), which represents a FDR≤0.1, was considered as a confident hit. The pathway enrichment analysis was performed using Metascape(50). The violin plots were generated using BoxPlotR (http://shiny.chemgrid.org/boxplotr/).

### Chemical Screen

OCI-AML2 cells were seeded in 25 μl per well in 384-well plates at the density of 300,000 cells/ml and treated with vehicle (0.1% DMSO) or compounds contained in the Anticancer compound library (MedChemExpress HY-L025) for 48 hours. A robot distributed 25 μl of the ATPlite luminescence assay reagent (ATPlite Luminescence Assay System; PerkinElmer) in each well. The contents were mixed for 2 minutes at 1100 rpm on an orbital shaker (OrbiShaker MP, Benchmark Scientific) and plates were incubated for 5 minutes at room temperature to stabilize luminescent signals. Units of luminescent signal generated by a thermo-stable luciferase are proportional to the amount of ATP presented in viable cells. Luminescence was recorded using a SpectraMax L384LW (Molecular Devices) reader with an acquisition time of 0.2s.

## Acknowledgements

This work was supported by the National Cancer Institute R35 CA210030 (K.S.), R50CA211404 (M.W.), the Children’s Leukemia Research Association (K.S.) and a St. Baldrick’s Foundation Robert J. Arceci Innovation Award (K.S). S.L. is a Fellow of the Leukemia and Lymphoma Society. B.K.A.S is supported by Department of Defense PRCRP Horizon Award (CA181249). N.V.D is supported by a St. Baldrick’s Foundation Fellowship. L.L. is a St. Baldrick’s Foundation Scholar. The Flow Cytometry Core at the Cincinnati Children’s Medical Center is supported by NIH S10OD023410. This work was also supported by the 4C’s Fund (K.S.).

## Author contributions

S.L. and C.L. conceived the study, designed and performed the experiments, analyzed the data, interpreted the results and wrote the manuscript. B.K.A.S assisted with the dTAG experiments, analyzed the data, and interpreted the results. A.R. and A.C. performed in vivo studies, analyzed the data, and interpreted the results. N.V.D., G.K and S.T.Y. designed the CRISPR library. N.V.D assisted with DepMap data analysis. S.T.Y assisted with the barcoding experiments and interpreted the results. M.K., C.W., S.M. and B.A. provided technical assistance, analyzed the data, and interpreted the results. T.N.M. and J.R. assisted with BH3-profiling experiments, analyzed the data, and interpreted the results. J.D.M. and A.L. provided resource support and intellectual input. F.P. assisted with screen data analysis. L.L. and M.W provided PDX resource and assisted with PDX-Cas9 model development. J.T. and K.S. conceived the study, designed the experiments, interpreted the results, supervised, and funded the study. All authors read, edited, and approved the final manuscript.

## Supplementary Figures

**Figure S1.**
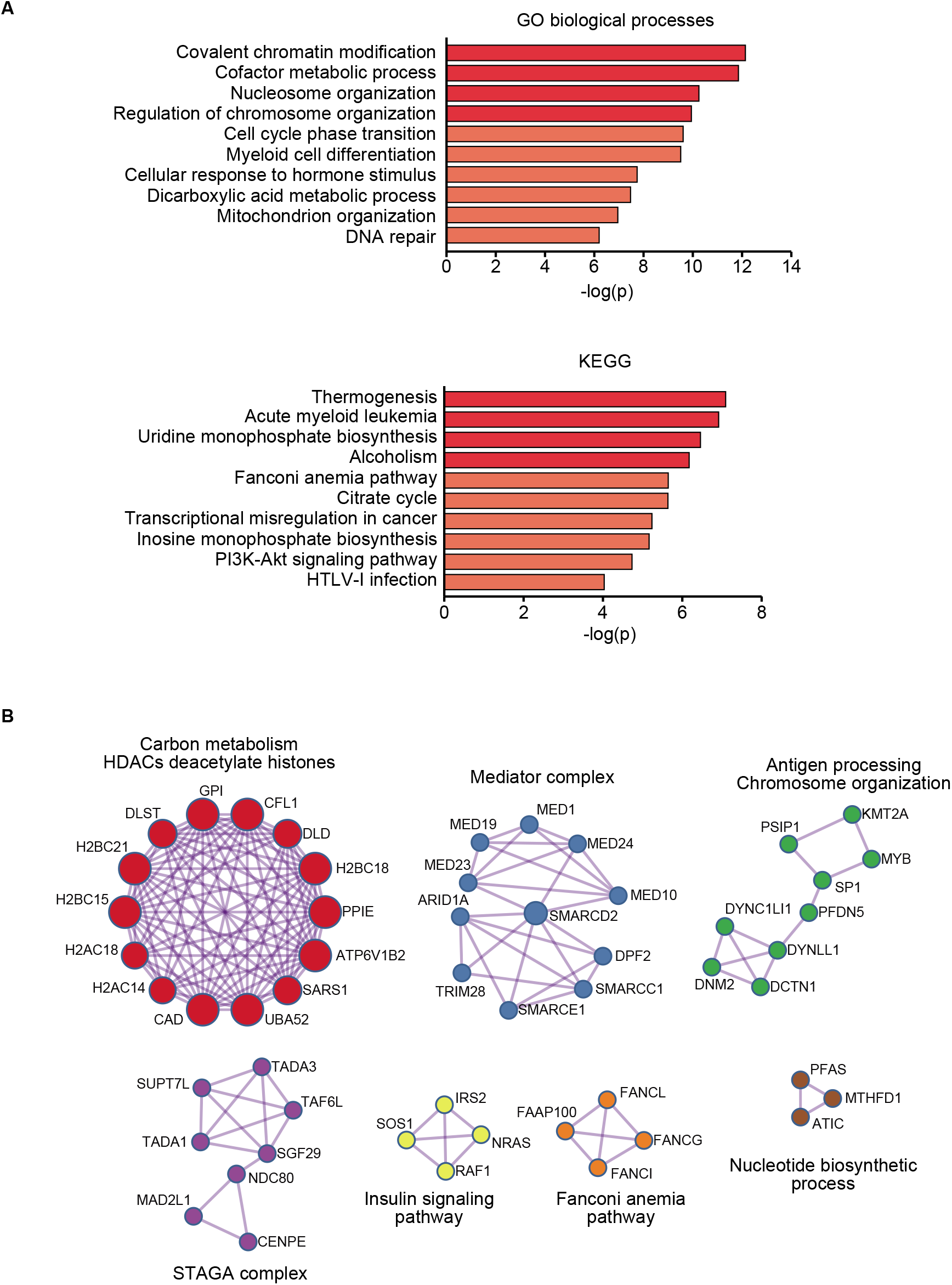
Pathway enrichment in the screen library. **A, B.** The enriched GO biological processes and KEGG pathways (**A**) and protein interaction network (**B**) in genes included in the library. The enrichment analysis was performed using Metascape(50).

**Figure S2.**
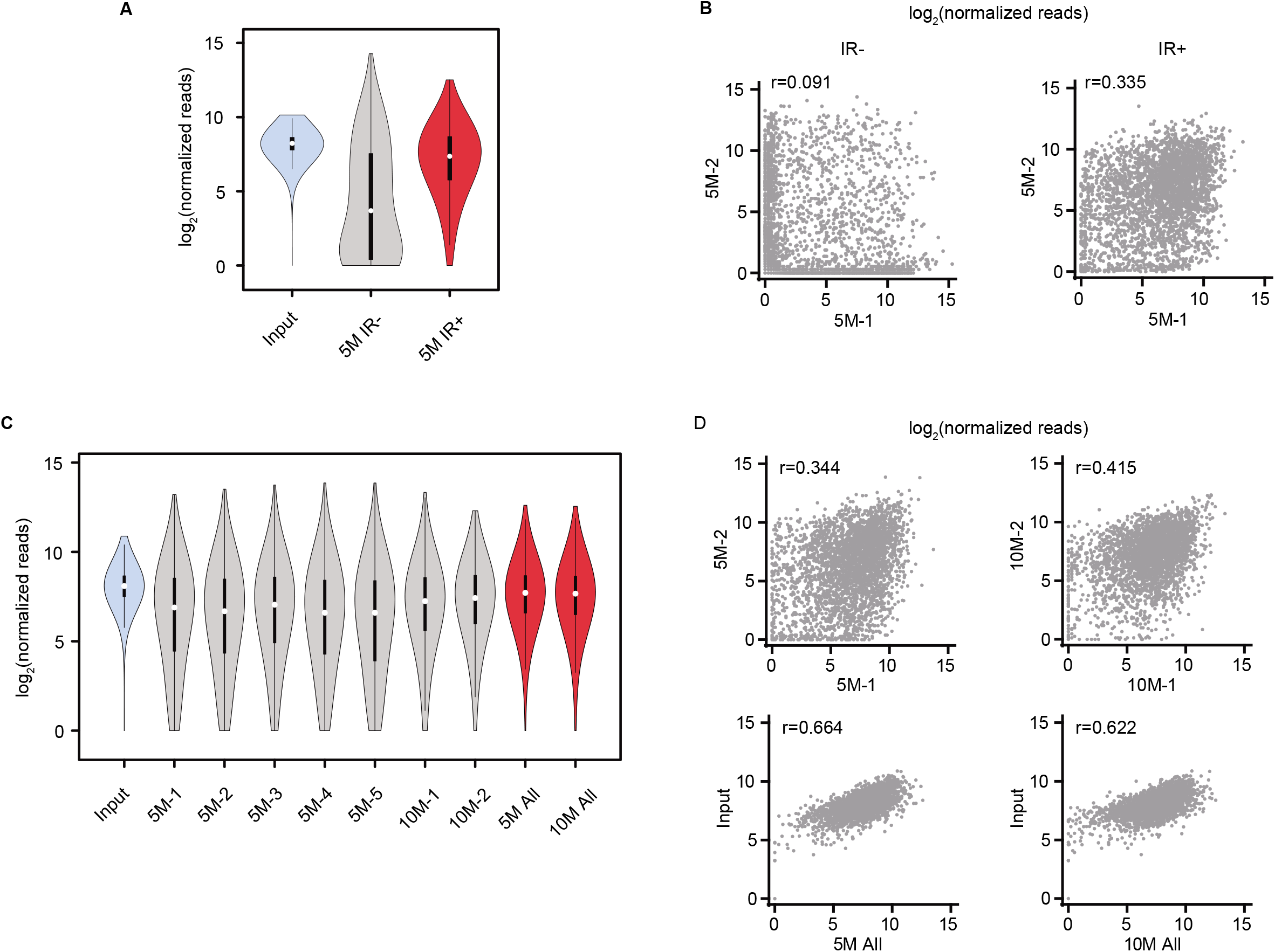
Barcoding experiments for screen optimization. **A.** Violin plot showing the distribution of barcodes in input references, mouse recipients without or with the sub-lethal irradiation (IR- and IR+, respectively). 5 million (5M) cells were transplanted into each mouse, bone marrow cells were analyzed, and normalized read counts from two mice of each group were averaged. White circles show the medians, box limits indicate the 25th and 75th percentiles, whiskers extend 1.5 times the interquartile range from the 25th and 75th percentiles, and polygons represent density estimates of data and extend to extreme values. **B.** Scatterplots showing the correlation of barcode read counts between two individual mice without or with sub-lethal irradiation (IR- and IR+, respectively). **C.** Violin plot showing the distribution of barcodes in the input references and the irradiated mouse recipients transplanted with 5 million (5M) or 10 million (10M) cells. 5M ALL and 10M ALL show the average normalized reads from 5 recipients of 5 million cells or 2 recipients of 10 million cells, respectively. **D.** Scatterplots showing the correlation of barcode read counts between two individual 5M or 10M recipients (upper), and the correlation between input and 5M ALL or 10M ALL(lower).

**Figure S3.**
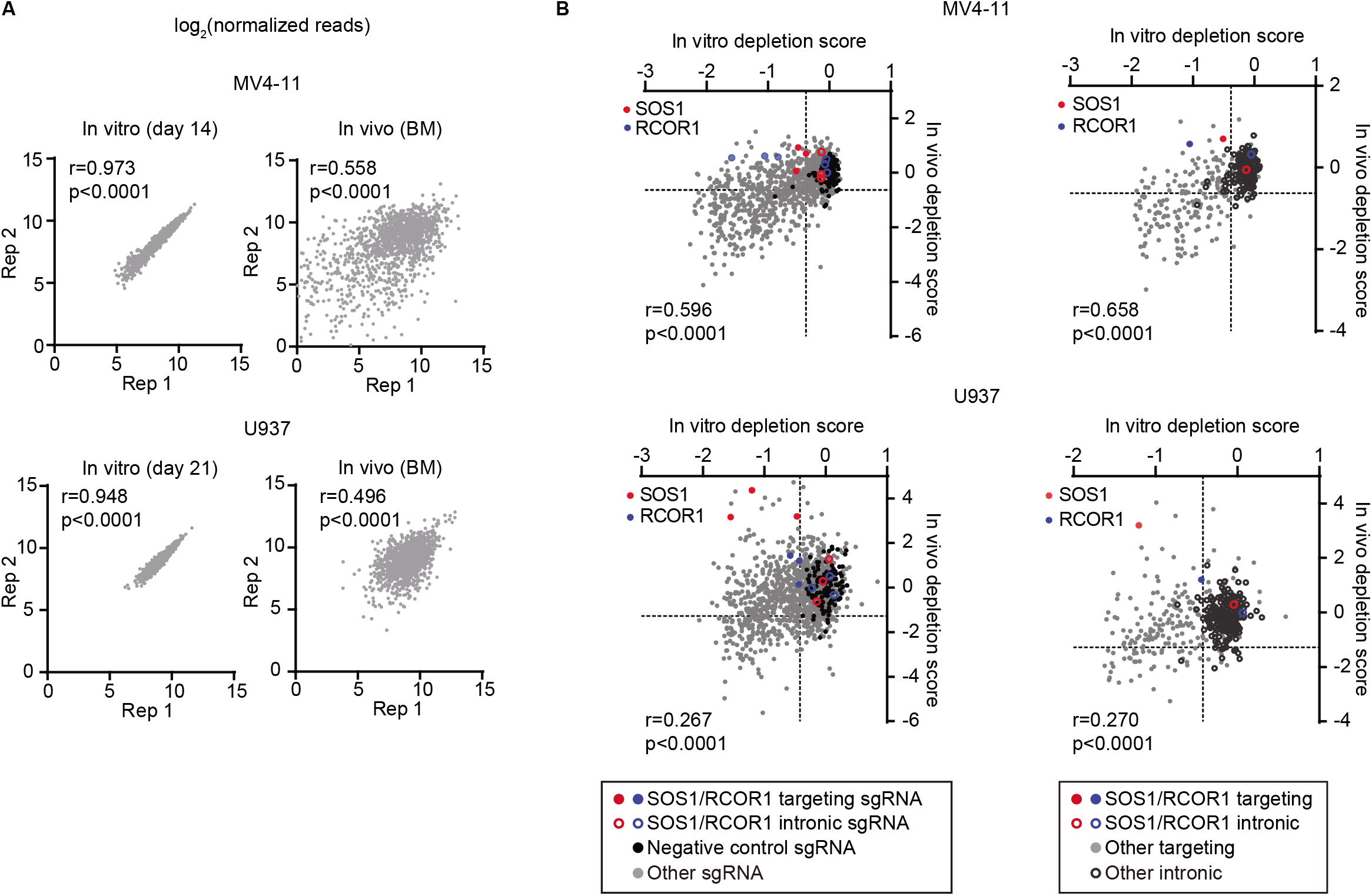
Identification of human AML dependencies in vivo. **A.** Scatterplots showing the correlation of sgRNA abundance between two replicates. **B.** Scatterplots showing the in vivo and in vitro depletion scores at sgRNA level (left panels) and gene level (right panels) in MV4-11 and U937 cells. Data points representing negative controls sgRNAs (black solid circles in left panels) and median value of each intronic sgRNA set (black hollow circles in right panels) are indicated. Scores of *SOS1* and *RCOR1* (red and blue solid circles) and their intronic controls (red and blue hollow circles) are highlighted as examples of the in vivo discrepancies from in vitro.

**Figure S4.**
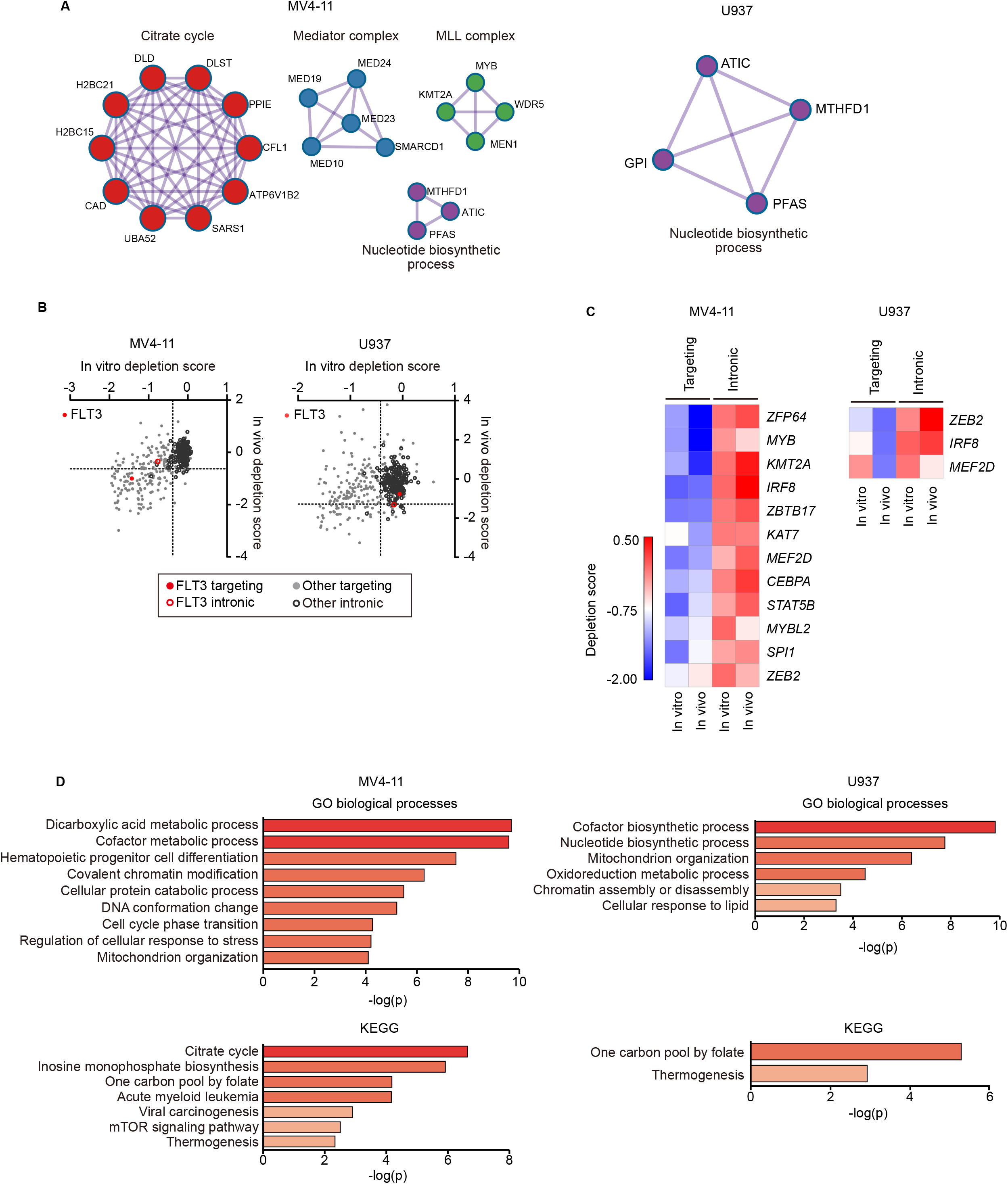
Pathways enriched in in vivo dependency. **A.** The enrichment of protein interaction network in genes scoring in vivo. **B.** Scatterplots showing the in vivo and in vitro depletion scores of *FLT3* (red solid circle) and its intronic controls (red hollow circle) at gene level in MV4-11 and U937. Data points representing the median value of each intronic sgRNA set are indicated (black hollow circles). **C.** Heatmap showing depletion scores of transcription factors as in vivo dependencies. Both targeting and intronic control scores are displayed. **D.** The enriched GO biological processes and KEGG pathways in genes scoring in vivo. The enrichment analysis in (**A**) and (**D**) was performed using Metascape.

**Figure S5.**
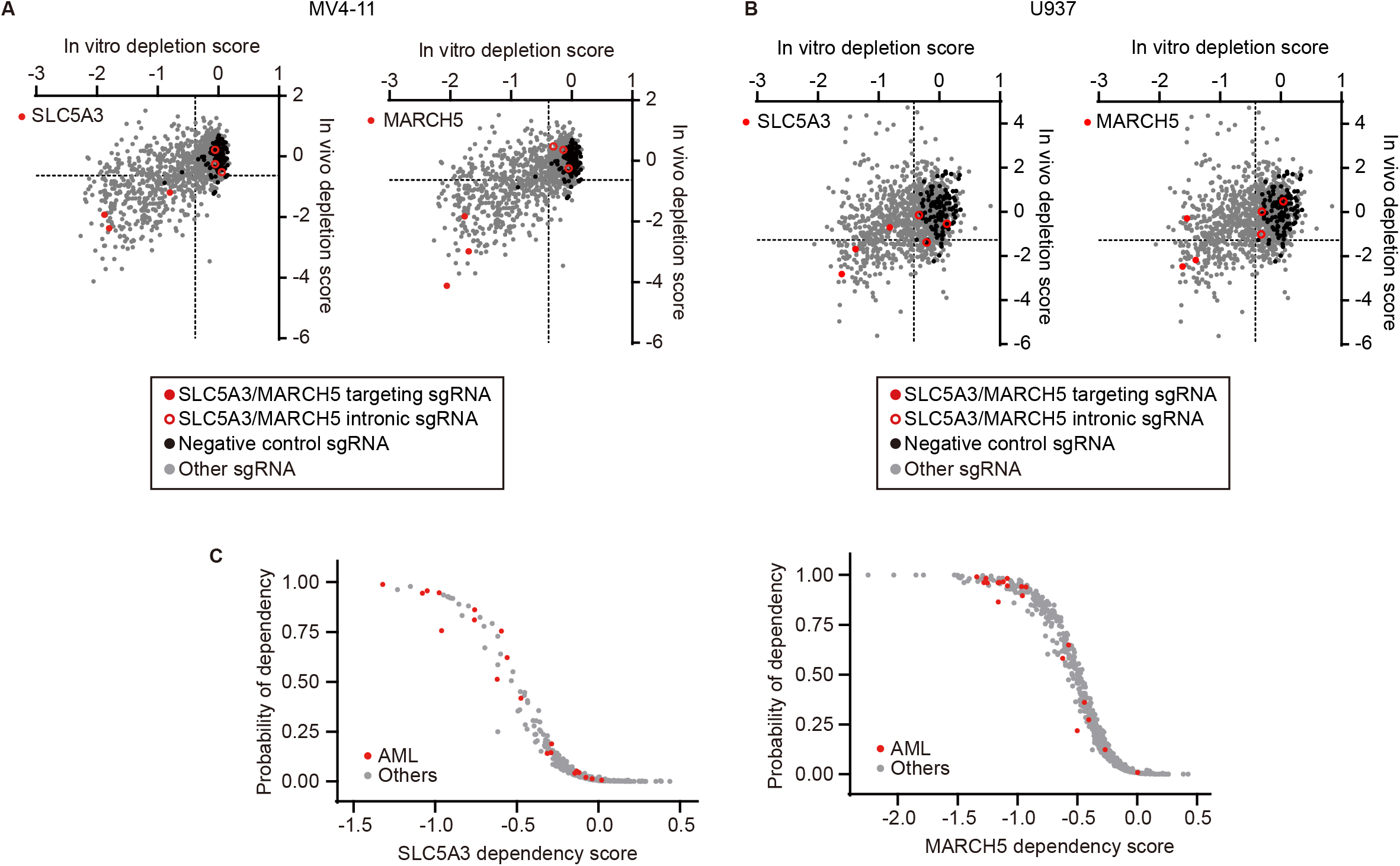
SLC5A3 and MARCH5 are strong AML dependencies. **A, B.** Scatterplots showing the in vivo and in vitro depletion scores of *SLC5A3* and *MARCH5* at sgRNA level in MV4-11 (**A**) and U937 (**B**). Data points representing negative controls sgRNAs (black solid circles) are indicated. Scores of *SLC5A3* and *MARCH5* targeting sgRNAs (red solid circle circles) and their intronic control sgRNAs (red hollow circle circles) are highlighted. **C.** Scatterplot of SLC5A3 and MARCH5 CERES dependency scores in expanded DepMap CRISPR dataset, 823 human cancer cell lines are included; each dot represents a cell line.

**Figure S6.**
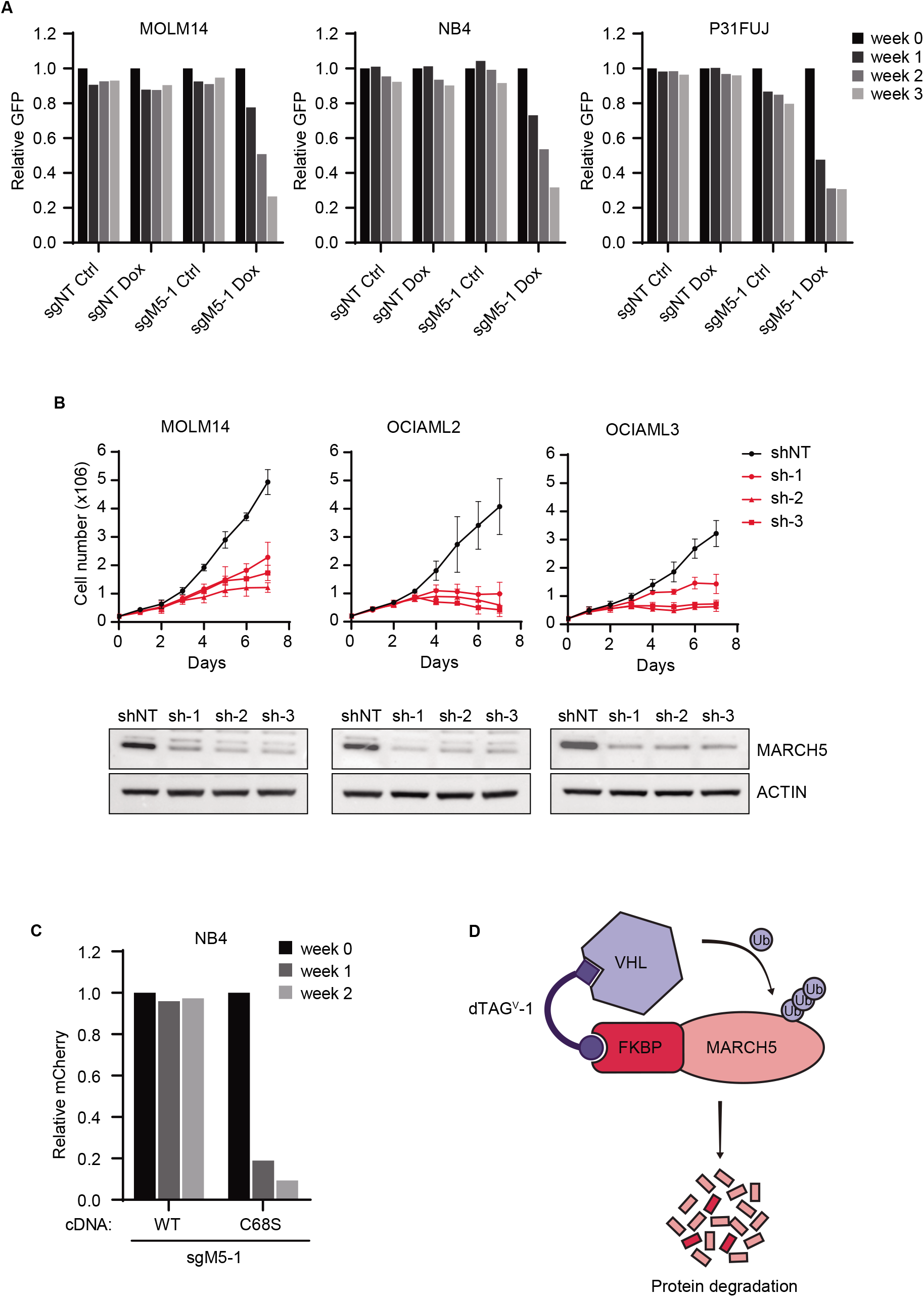
MARCH5 depletion inhibits AML growth. **A.** Competition proliferation assay to evaluate growth of AML cell lines transduced with Dox-inducible sgNT or *MARCH5* sg-1 (sgM5-1). **B.** Cumulative cell numbers of AML cell lines transduced with Dox-inducible control (shNT) or MARCH5-targeting shRNAs (sh-1~3). n=3, results represent mean ± SD. **C.** Competition proliferation assay of NB4 cells overexpressing wildtype *MARCH5* (WT) or ligase defective mutant (C68S) and transduced with mCherry-linked *MARCH5* sgRNA. **D.** Schematic of the dTAG system for controlled protein degradation.

**Figure S7.**
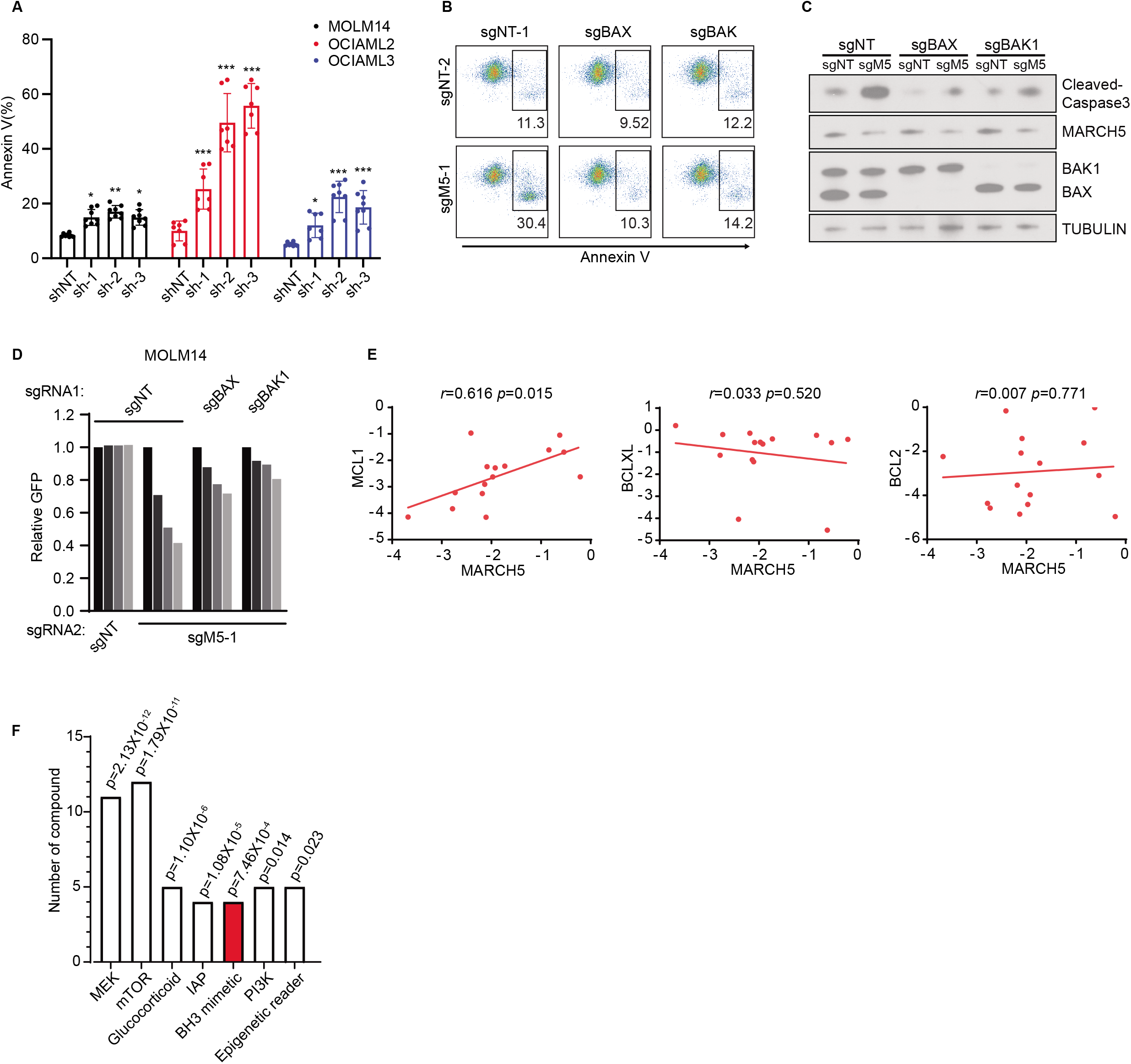
MARCH5 depletion induces apoptosis. **A.** Percentage of Annexin V analyzed by flow cytometry in AML cells transduced with the indicated shRNAs. *p<0.05, **p<0.01, ***p<0.001, p values were calculated by unpaired two-tailed t-test. **B, C.** Flow cytometry analysis of Annexin V (**B**) and immunoblot analysis of cleaved-caspase3 (**C**) in control, *BAX-* and *BAK1-* knockout MV4-11 cells transduced with *MARCH5* sgRNA. **D.** Competitive growth of control, *BAX-* and *BAK1-* knockout MOLM14 cells transduced with *MARCH5* sgRNA. **E.** Scatterplots showing the correlation of dependency scores between *MARCH5* and *MCL1, BCL2* or *BCLXL* in AML CRISPR screen from Wang & Sabatini et al. (13). **F.** Categories of targets enriched in sensitizing compounds. p values were calculated by a hypergeometric test compared to the composition of the screen library.

**Figure S8.**
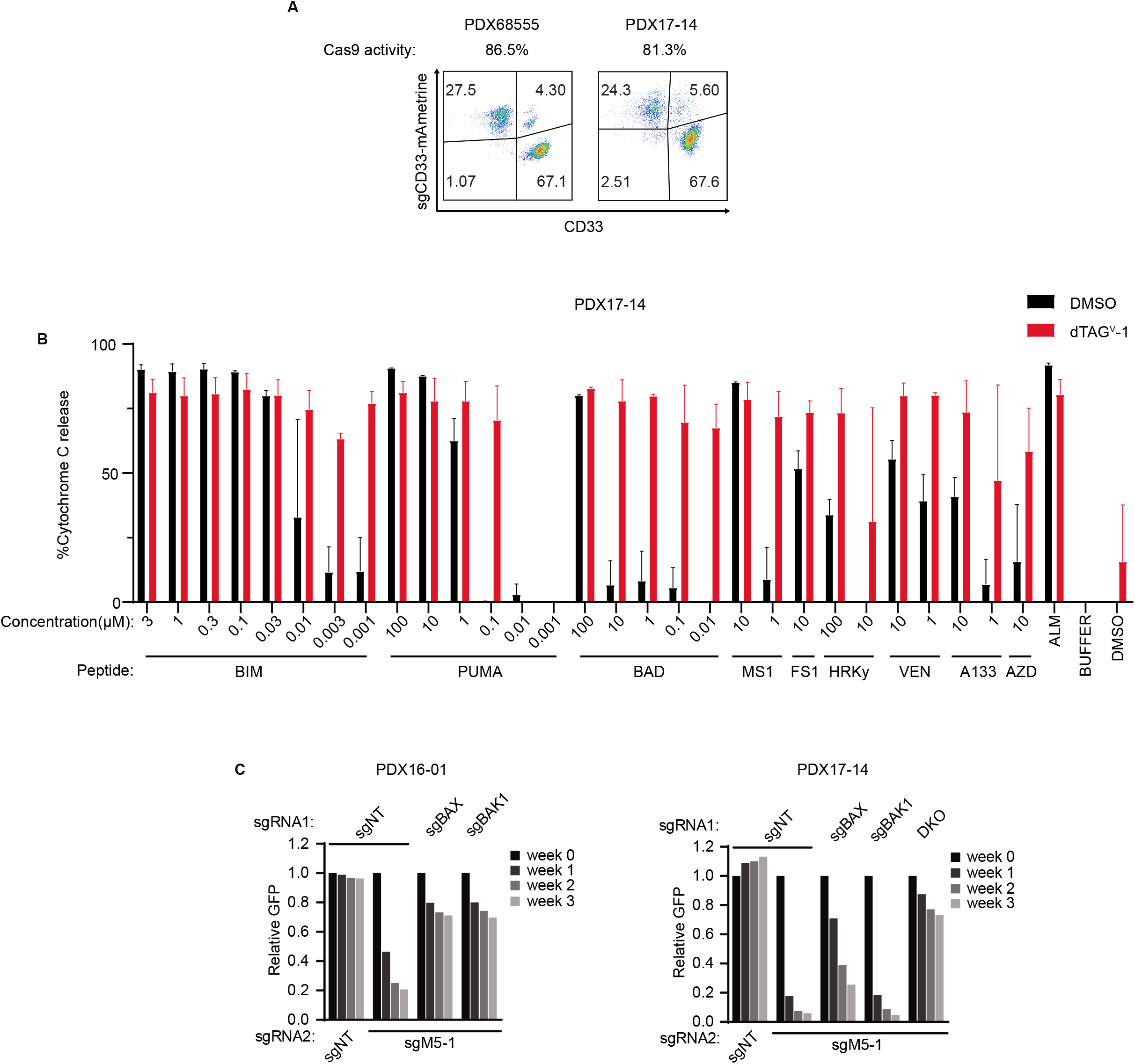
Evaluating AML dependency targets in Cas9-competent PDX models. **A.** Flow cytometry analysis to evaluate Cas9 activity in Cas9-expressing PDX models using CD33 expression as a reporter. PDX cells were transduced with mAmetrine-linked sgRNA targeting *CD33,* the percentage of Cas9 active cells was indicated by the ratio of CD33-mAmetrine+ cells to total mAmetrine+ cells. **B.** BH3 profiling to measure the apoptotic priming of MARCH5-dTAG PDX17-14 cells treated with DMSO or 500 nM dTAG^V^-1 for 2 hours. MS1 is a peptide targeting MCL1; FS1 is a peptide targeting BCL2A1; VEN is venetoclax; A133 (A1331852) is a BCLXL inhibitor; AZD (AZD5991) is a MCL1 inhibitor; ALM (alamethicin) serves as a positive control for cytochrome C release. Results represent mean ± SD, n=2. **C.** Competition proliferation assay to evaluate the growth effect of *MARCH5* knockout in control, *BAX-, BAK1-* and double(DKO)-knockout PDX cells.

